# Acyltransferases in the first step of glycerophospholipid synthesis have redundant and non-redundant roles in *sn-*1 acyl chain regulation, ether lipid levels, and cell survival

**DOI:** 10.1101/2024.01.17.576130

**Authors:** Sarah Salame, Hasna Qasmaoui, Gregor P Jose, Lucile Fleuriot, Delphine Debayle, Takeshi Harayama

## Abstract

The increased use of lipidomics analyses in biomedical research made it crucial to understand correctly how lipid compositions are regulated. Lipid acyl chains are important regulators of membrane physicochemical properties, and are incorporated by various acyltransferases. The acyltransferases implicated in the first step of glycerophospholipid synthesis, the GPATs and GNPAT, have distinct localizations and substrate preferences. This suggests that they have distinct roles on lipid regulation, but a complete understanding of their redundant and non-redundant functions is missing. We report here a comprehensive analysis of cells having mutations in GPATs and GNPAT, either alone or in combinations. Our results suggest that the balance between GPATs and GNPAT affect the levels of ether lipids, together with glycerophospholipid acyl chain length and unsaturation. In addition, we found that lipid synthesis initiated at peroxisomes, but not mitochondria, is sufficient to provide the lipid synthesis flux required for normal cell growth. Our study unveils the multifaceted roles of the first step of de novo glycerophospholipid synthesis, thus leading to a better understanding of how lipidomes are shaped.

## 1. Introduction

Membrane lipid structures have influences on membrane physicochemical properties, membrane-associated protein activity and localization, and metabolic fates of lipid themselves^1,2^. Lipid acyl chain structures (double bond numbers and positions or length) are important determinants of membrane fluidity, flexibility, and impermeability^3^. Glycerophospholipids (GPLs) have typically an asymmetric *sn-*1 saturated/monounsaturated *sn-* 2 mono/polyunsaturated configuration of acyl chains, albeit with multiple exceptions^3^. The maintenance of this asymmetry is fundamental for life, as the overaccumulation of di-saturated GPLs triggers endoplasmic reticulum (ER) stress, while membranes made from di-polyunsaturated GPLs become highly permeable^3–6^. Therefore, the proper regulation of GPL acyl chains at both *sn-*1 and *sn-*2 positions is critical to maintain cellular functions.

Acyl chains are incorporated in GPLs by acyltransferases^7,8^. Glycerol-3-phosphate *O-* acyltransferases (GPATs) and glyceronephosphate *O-*acyltransferase (GNPAT) catalyze the first step of GPL synthesis^8^ (see Figure 1 for metabolic pathways described in this paragraph). GPATs use glycerol-3-phosphate (G3P) and acyl-CoA to synthesize 1-acyl lysophosphatidic acid (acyl-LPA), the precursor of GPLs and triglycerides (TAGs) having ester-linked acyl-chains at the *sn-*1 position. There are four known mammalian GPATs, with GPAT1 (called GPAM in this study, based on its official gene name) and GPAT2 being expressed in mitochondria, while GPAT3 and GPAT4 are found in the ER^9^. GNPAT is expressed in peroxisomes, and uses dihydroxyacetone phosphate (DHAP) and acyl-CoA to synthesize acyl-DHAP^10^. Acyl-DHAP is either converted into acyl-LPA or further metabolized into alkyl-DHAP and 1-alkyl lysophosphatidic acid (alkyl-LPA). Alkyl-LPA is used in the ER for the synthesis GPLs and TAGs with ether-linked *sn-*1 alkyl chains, namely ether-GPLs and ether-TAGs^10^. Therefore, GPL biosynthesis is initiated by GPATs and GNPAT at various organelles (ER, mitochondria, and peroxisomes). However, the biological significance of this compartmentalization in lipid synthesis remains unclear. Some studies suggested that the lack of lipid synthesis initiation in an organelle can be compensated by other organelles. For example, in *Caenorhabditis elegans*, the loss of GPATs is lethal only when ER GPATs and mitochondrial GPAT are all mutated^11^. In mammalian cells, the combinatory loss of GNPAT and CHP1, the activator of GPAT4, inhibits cell proliferation^8^. These studies suggest that there is redundancy in lipid synthesis even when it is initiated in distinct organelles. However, it remains unclear to which extent lipid synthesis at individual organelles can compensate the loss of lipid synthesis initiation in other organelles.

**Figure 1.**
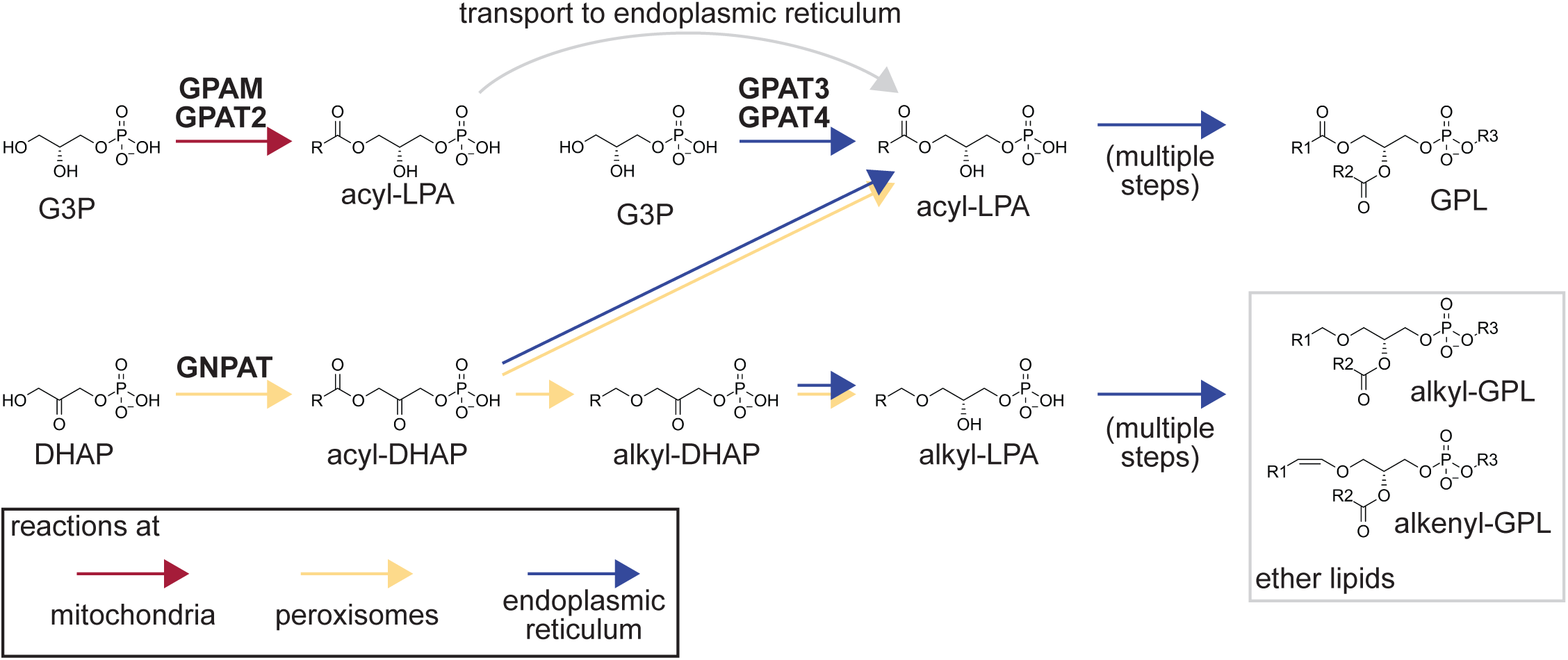
Metabolic pathways analyzed in this study. The first steps of GPL *de novo* synthesis are illustrated, with the localization of reactions color coded. The reactions with two different arrows can take place in multiple organelles.

Besides their localization, GPATs have differences in acyl-CoA preferences, at least when analyzed with *in vitro* assays^12^. Given the importance of proper GPL acyl chain compositions in cells, it is critical to know whether acyl-CoA preferences of GPATs and GNPAT affect lipid composition in cellular contexts. GPAM knockout mice have reductions in GPLs with palmitic acid (16:0) in heart^13^. GPAT4 is proposed to favor the synthesis of acyl-LPA with stearic acid^14^ (18:0). However, in most cases, the contribution of GPATs and GNPAT on acyl chain regulation have been analyzed separately, in different cells or tissues, and in different contexts. Therefore, a side-by-side analysis of how these enzymes regulate acyl chains would be valuable for the correct understanding of lipid homeostasis. In addition, given the non-redundant role of GNPAT on ether lipid biosynthesis, it is crucial to know whether the balance between GPATs and GNPAT affect ether lipid levels, and to which extent GNPAT can contribute to non-ether lipid synthesis.

In this study, we generated multiple mutant cell lines lacking GPATs and/or GNPAT, either alone or in combination, to analyze the ability of cells to survive and generate ether and ester lipids with distinct acyl chains.

## 2. Materials and Methods

### Cell culture

HeLa cells (HeLa MZ clones kindly provided by Jean Gruenberg, University of Geneva) were cultured in Dulbecco’s Modified Eagle Medium (high glucose DMEM, GlutaMAX, pyruvate, gibco) supplemented with 10% fetal calf serum (eurobio scientific) and 1% ZellShield cell culture contamination preventive (Minerva-Biolabs). Cells were passaged at 80-90% confluency, by washing them using Dulbecco’s Phosphate-Buffered Saline (DPBS without calcium/magnesium, gibco) two times, followed by their detachment using trypsin-EDTA (gibco). Trypsinization was stopped by adding complete medium and passages were performed in the range of 1:5 to 1:20 dilutions. Cell counting was done using Countess II automated Cell Counter (invitrogen) according to manufacturer instructions.

### Generation of mutant cell lines

Mutant cell lines were generated by the CRISPR-Cas9 strategy GENF (gene co-targeting with non-efficient conditions), which is highly efficient and does not necessitate clone isolation^15^. Mutations are introduced in target genes together with the HPRT1 gene, which confers resistance against 6-thioguanine and allows drug selection of HPRT1 mutant cells. By introducing HPRT1 mutations using a low efficient sgRNA and selecting HPRT1 mutant cells with 6-thioguanine, we isolate cells with high CRISPR-Cas9 efficiency, thus strongly enriching target mutated cells. The sgRNA target sequences used in this study are listed in supplemental file 1.

HeLa cells (80000 cells/well) were reverse transfected in 24 well plates with 495 ng pX330-based plasmids (encoding Cas9 and target sgRNAs, deposited by Feng Zhang as addgene plasmid #42230) and 5 ng pHPRTsg (encoding HPRT1 sgRNA, target sequence CGATAATCCAAAGATGGTCA, which has low efficiency due two mismatches at the region distal-most from the protospacer adjacent motif) using Lipofectamine 3000 (invitrogen). 5 days post-transfection, cells were selected with 6 µg/mL 6-thioguanine (Sigma-Aldrich). The generated mutant cells were stocked in CELLBANKER 1 (Nippon Zenyaku Kogyo).

Mutation rates were analyzed by PCR amplification of target regions from genomic DNA, Sanger sequencing of the product, and its analysis by a decomposition software (see below). Genomic DNA was isolated by lysing cells (10 mM Tris-HCl pH 8.0, 1 mM EDTA, 0.67% SDS, and 125 μg/ml proteinase K from Roche), followed by incubation 4 hours at 55°C with weak agitation. Genomic DNA was precipitated by isopropanol in the presence of 500 mM NaCl. PCR was done using ExTaq or PrimeSTAR GXL, with primers listed in supplemental file 1. PCR products were treated with FastAP alkaline phosphatase and Exonuclease I (Thermo Scientific), and Sanger sequencing was performed on a 3130xl genetic analyzer (ABI), using the forward primers used for PCR.

TIDE (Tracking of Indels by Decomposition) was used to analyze indel patterns in mutated cells^16^. It decomposes indel patterns from Sanger sequenced PCR amplicons, by comparing data obtained from mutant cells and control cells. From the results of TIDE, the loss of wild type sequence (indel = 0) was used as a measure of successful gene mutation, thus mutation rates are illustrated as 100 - %wild type sequence. We validated this approach as being more representative of high mutation rates in a previous study^15^. The sum of mutations having in-frame indels (indel size = 3N) was calculated and divided by the sum of all obtained indels to analyze the percentage of in-frame mutations that potentially generate functional proteins.

At the end of 6-thioguanine selection, mutant cells were detached and reseeded into 24 well plates with a serial dilution. After attachment, cells were fixed with methanol and stained with Crystal violet to investigate the lethality of mutants.

### Lipidomics analysis

Samples were prepared by seeding 750,000 cells on 6 cm dishes. 24 hours later, cells were washed and scraped in ice-cold PBS, centrifuged at 2,500 rpm at 4°C, and pellets were snap frozen in liquid N_2_.

For lipid extraction, pellets were resuspended in 200 µL water, and 500 µL of methanol was added. Internal standards (4 µL of SPLASH LIPIDOMIX Mass Spec Standard and 20 pmol each of C17 ceramide, C17 Glucosylceramide, and C17 Lactosylceramide, Avanti Polar Lipids) were added to the mixture, followed by 250 µL of chloroform, and the single phase was shaken for 10 minutes. Then, 250 µL of water and 250 µL of chloroform were added, and the mixture was shaken for 10 minutes. After centrifugation for 15 minutes at 3,000 rpm, 400 µL of lower phase were transferred to new glass tubes and evaporated under nitrogen. Extracts were resuspended in 60 µL isopropanol : methanol : water (5:3:2) and 3 µL was injected into a Ultimate 3000 UHPLC system coupled to a Q Exactive mass spectrometer (Thermo Fisher Scientific). Separation was done on an Accucore C18 column (150 x 2.1 mm, 2.6 µm particles), using a gradient of solvent B (isopropanol : acetonitrile : water (88:10:2, v/v) supplemented with 2 mM ammonium formate and 0.02% formate) over solvent A (acetonitrile : water (1:1, v/v) supplemented with 10 mM ammonium formate and 0.1% formate) at a flow rate of 400 µL/minute. The gradient started at 35% solvent B and was changed linearly to reach 60% B at 4 minutes, 70% B at 8 minutes, 85% B at 16 minutes, 97% B at 25 minutes, and 100% B at 25.1 minutes. 100% B was maintained until 31 minutes, and the column was reconditioned at 35% B for 4 minutes. Separate injections were done for analysis in positive and negative ion modes. Data was acquired by data dependent MS2. MS1 was analyzed at a resolution of 70,000 at *m/z* = 200 on the range of *m/z* = 250-1200 with an AGC target of 1000000 and maximum injection time of 250 msec. For MS2, 15 precursor ions were selected in an isolation window of *m/z* = 0.95 and fragmented by higher-energy collisional dissociation with a normalized collision energy of 25 eV and 30 eV for positive mode and 20 eV, 30 eV, and 40 eV for negative mode, and injected into the Orbitrap analyzer with an AGC target of 100000 and maximum injection time of 80 msec, and analyzed at a resolution of 35,000 at *m/z* = 200.

LC-MS data were analyzed using MS-DIAL5^17^. Peak interpretations were curated based on retention times and investigation of fragments corresponding to co-eluting peaks. Annotation of ether and vinyl ether lipids was done based on retention time^18^, and was confirmed by whether the peaks disappear in mutant cells lacking PEDS1 (GPJ, unpublished data), the enzyme responsible for vinyl ether bond formation in GPLs. Lipids were semi-quantified by dividing peak areas with those of internal standards and multiplying with the calculated quantities added to samples. Lipid levels were normalized with the sum of phosphatidylcholine, phosphatidylethanolamine, phosphatidylinositol, and phosphatidylserine, as % of total GPLs. All lipidomics data of this study are provided in supplemental file 2.

### Data processing

The GPLs were identified in the form of XX (lipid class) FA1_FA2, where FA1 is the more saturated acyl chain and the shorter one if the saturation is the same as FA2. We summed the quantities of all the lipids from the same species and the same acyl chain found either as FA1 or as FA2. When both FA1 and FA2 were identical (e.g. PC 16:0_16:0), the quantities were summed two times. For example, for the analysis of 16:0 in PC, we summed PC 14:0_16:0, PC 16:0_16:0 (two times), PC 16:0_18:1, etc. The same calculation was done for all detected acyl chains, and everything was summed up to obtain total values. Then individual values for each acyl chain were divided with this total to obtain percentages. The same calculations were done for all four GPL classes analyzed in this study. Then, we repeated the whole process while considering only FA1, which is more likely to reflect *sn-*1 acyl chains, which are the major targets of this study.

## 3. Results

### Generation of GPAT/GNPAT mutant cells and analysis of lipid classes

We mutated individual GPATs or GNPAT in HeLa cells to perform lipidomics and analyze their contributions on ether lipid levels and GPL acyl chains. We used a CRISPR-Cas9 strategy GENF (gene co-targeting with non-efficient conditions) developed recently to mutate targets in a polyclonal cell population, which is beneficial for lipidomics analyses due to reduced artifacts linked to clone isolation^15^. All targets were mutated with >95% mutation rates (100% for GPATs, namely GPAM, GPAT3, and GPAT4, Figure 2A). GPAT2 was not included in our analysis, as its expression is very low in HeLa cells (Figure S1).

**Figure 2.**
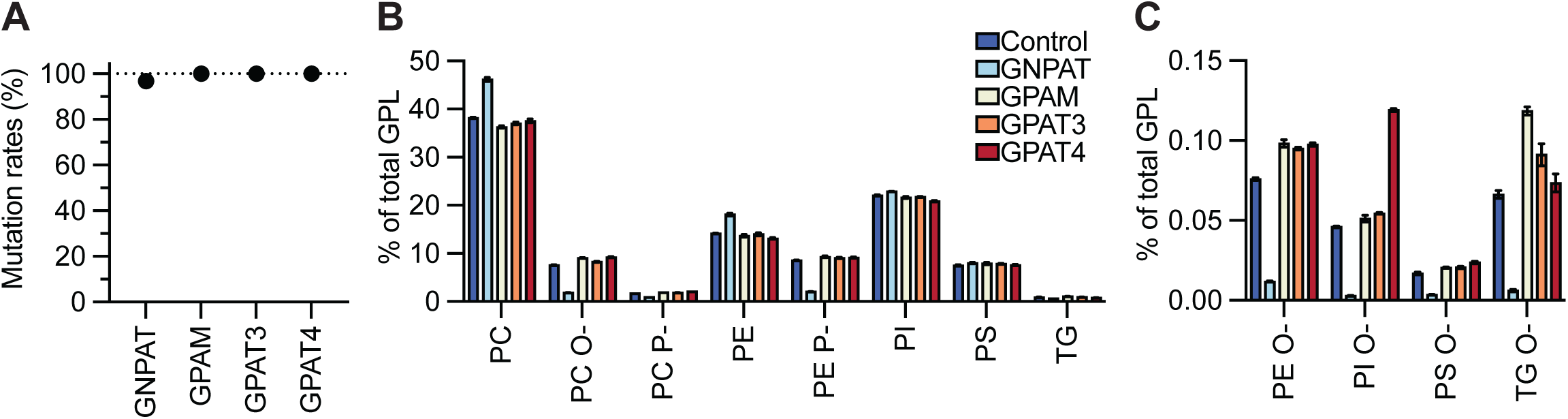
Analysis of single GPAT/GNPAT mutants. (A) Mutation rates in the obtained polyclonal mutants analyzed by TIDE. (BC) Analysis of lipid changes in mutants of the indicated genes, at the lipid class/subclass levels. Error bars are SEM (n=4). Results of statistical tests are provided in a supplemental file 3. See also Figure S1 for the expression levels of target genes.

Using these mutants, we performed untargeted lipidomics and measured the levels of phosphatidylcholine (PC), phosphatidylethanolamine (PE), phosphatidylinositol (PI), phosphatidylserine (PS), and triglycerides (TG). We analyzed ether species too (labeled O-for alkyl lipids and P-for alkenyl lipids, for TGs we labeled them as O-, but discrimination is not done), including minor ether PI and PS species (Figure 2BC, see y axis values). At lipid class levels, GPAT mutants did not have large changes in lipid composition, while GNPAT mutants had drastic decreases in ether GPLs and ether TG (Figure 2BC). GPAT mutants tended to have increased ether GPLs, which ranged between 5-25%, depending on the ether GPL class and the mutated GPAT (Figures 2BC). The increase in ether PI was exceptionally high in GPAT4 mutants (Figure 2C). The results suggested that GPATs and GNPAT compete for lipid synthesis, and that ether lipid levels increase proportionally to the contribution of GNPAT. Minor ether PI was especially sensitive to this trend.

### Global analysis of acyl chain dysregulation in GPAT/GNPAT mutants

Using the whole lipidomics dataset of individual lipid species, we performed principal component analysis (PCA) to reduce the dimensions of the data and visualize how mutants differ from control cells (Figure 3A). Only GNPAT mutants were discriminated with the first principal component (PC1), suggesting that the analysis is strongly influenced by the large changes in ether lipids induced by GNPAT loss.

**Figure 3.**
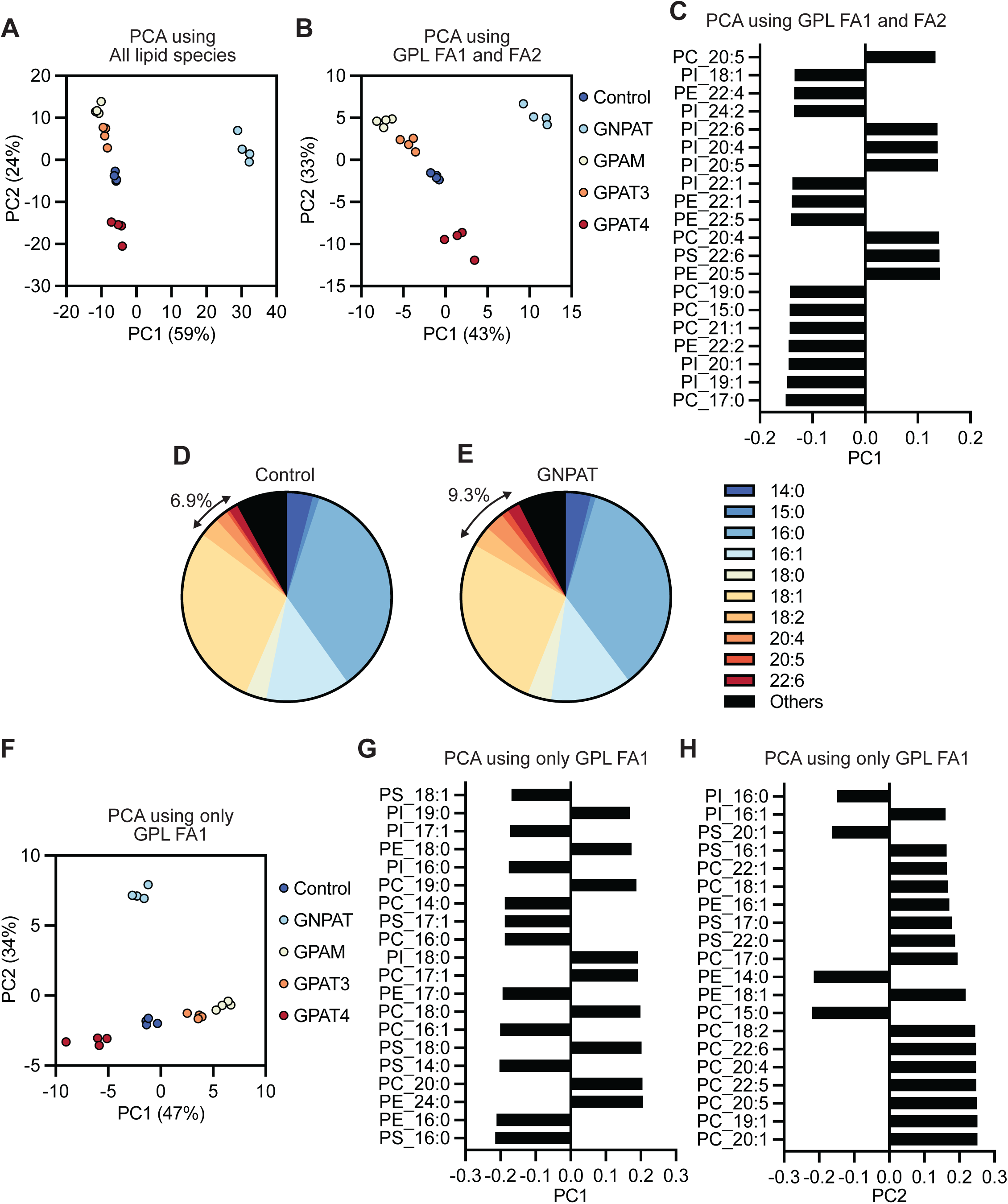
Multivariate analyses of GPAT/GNPAT mutants. (AB) Results of PCA depending on the data used as an input. All lipid species were used for (A), while the percentages of acyl chains that constitute FA1 and FA2 in each GPL class were used for (B). (C) PC1 loading for the PCA of (B). Note the abundance of PUFAs. (DE) Percentage of acyl chains constituting FA1 and FA2 in PC in the control and GNPAT mutants. Only species with >1% abundance in either cell line are shown, while “Others” illustrate the minor species and the species for which unequivocal acyl chain annotation could not be done by LC-MS. (F) Result of PCA, for which the input data was the percentage of acyl chains that constitute FA1 in each GPL class. (GH) PC1 and PC2 loadings for the PCA of (F). See also Figure S2 for the acyl chain profiles of control cells, obtained after different strategies of data simplification.

Therefore, we used an approach to simplify our dataset and give more focus on acyl chains, as briefly explained here (see materials and methods for more details). Our lipidomics approach allows discrimination of acyl chains when isomeric lipids are chromatographically separated, but does not provide information about *sn-*1/2 positions. Thus, acyl chains are illustrated as FA1_FA2, which does not specify positions (for example, PC 16:0_20:4 can be 1-16:0/2-20:4 or 1-20:4/2-16:0). FA1 is the more saturated and shorter (if same saturation as FA2) acyl chain, which is typically the acyl chain found at the *sn-*1 position, due to the saturated/unsaturated asymmetry between *sn-*1/2 positions^3^. To simplify the dataset with more focus on GPL acyl chains, we summed all (non-ether) GPL species having a common acyl chain irrespectively of FA1 or FA2, calculated their percentage within their classes. This gave information about which acyl chains are present in each lipid class, independently on their combinations (see control cell profiles in Figure S2A-D). Using this dataset, we repeated PCA, which discriminated GPAT mutants with PC1 better than the previous analysis (Figure 3B). However, GNPAT remained the most divergent mutant. Analysis of PCA loadings revealed which acyl chains contributed most to PC1, where we detected multiple polyunsaturated species and a few minor odd-chain species (Figure 3C). This suggested that our analysis was more influenced by secondary changes in polyunsaturated fatty acids (PUFAs) seen under ether lipid deficiency^19^, rather than changes in *sn-*1 acyl chains incorporated by GPATs/GNPAT. Indeed, examination of the extracted data of acyl chains revealed that GNPAT mutants had more PUFAs than control cells (Figure 3DE).

To let us have more focus on *sn-*1 acyl chains, we repeated the same data simplification process, but this time analyzing only FA1 (which is more likely the typical *sn-*1 acyl chain, as described above, see control cell profiles in Figure S2E-H). With this procedure, PCA revealed clear differences in GPAT mutants at the level of PC1 (Figure 3F). We saw that the lipidome of GPAT4 mutants change in opposite directions from GPAT3 and GPAM mutants. GPAT3 mutants changed in the same direction as GPAM mutants, but in a weaker manner, suggesting that GPAT3 and GPAM have some redundancy in acyl chain regulation. GNPAT mutants were discriminated from controls only at the level of PC2. This showed the benefit of simplifying lipidomics datasets depending on the biological question, even when the lipidomics procedure would not allow it in a strict manner. PCA loadings revealed a clear pattern, with saturated acyl chains changing in opposite directions depending on the chain length, with C17 being the threshold (Figure 3G). Thus, GPAT4 and GPAT3/GPAM seem to regulate the ratio between 16:0 and 18:0 acyl chains. Loading analysis of PC2, the major discriminant of GNPAT mutants, still detected PUFA changes (Figure 3H). Nevertheless, we found one odd chain species (15:0 in PC) that contributes to PC2.

### Regulation of acyl chains by GPATs/GNPAT

Guided by the results of PCA, we investigated how mutations in GPATs/GNPAT affected 16:0 and 18:0 as FA1 of various GPL classes. GPAM mutation reduced 16:0 levels in all analyzed GPLs (Figure 4A-D), while the effect of GPAT3 mutation was less pronounced. GPAT4 mutants had opposite changes, with 16:0 increases in all analyzed GPLs, except PC (Figure 4A-D). The levels of 18:0 decreased in PC, PS, and PI of GPAT4 mutants, while they were unchanged in PE (Figure 4E-H). We also analyzed shorter or longer acyl chains, and consistently with what PCA suggested, 14:0 levels of PS decreased in GPAM mutants and increased in GPAT4 mutants (Figure 4I). The changes in GPAT3 mutants were not significant. In contrary, 24:0 levels of PE increased in GPAM mutants and decreased in GPAT4 mutants (Figure 4J). This confirmed the opposite effects on saturated acyl chain length regulation by GPAM and GPAT4, which favor acyl chain shorter and longer than C17, respectively. Changes in 18:1 levels depended on the GPL class, with the effects of GPAM or GPAT4 mutations not being necessarily opposite (Figure 4KL).

**Figure 4.**
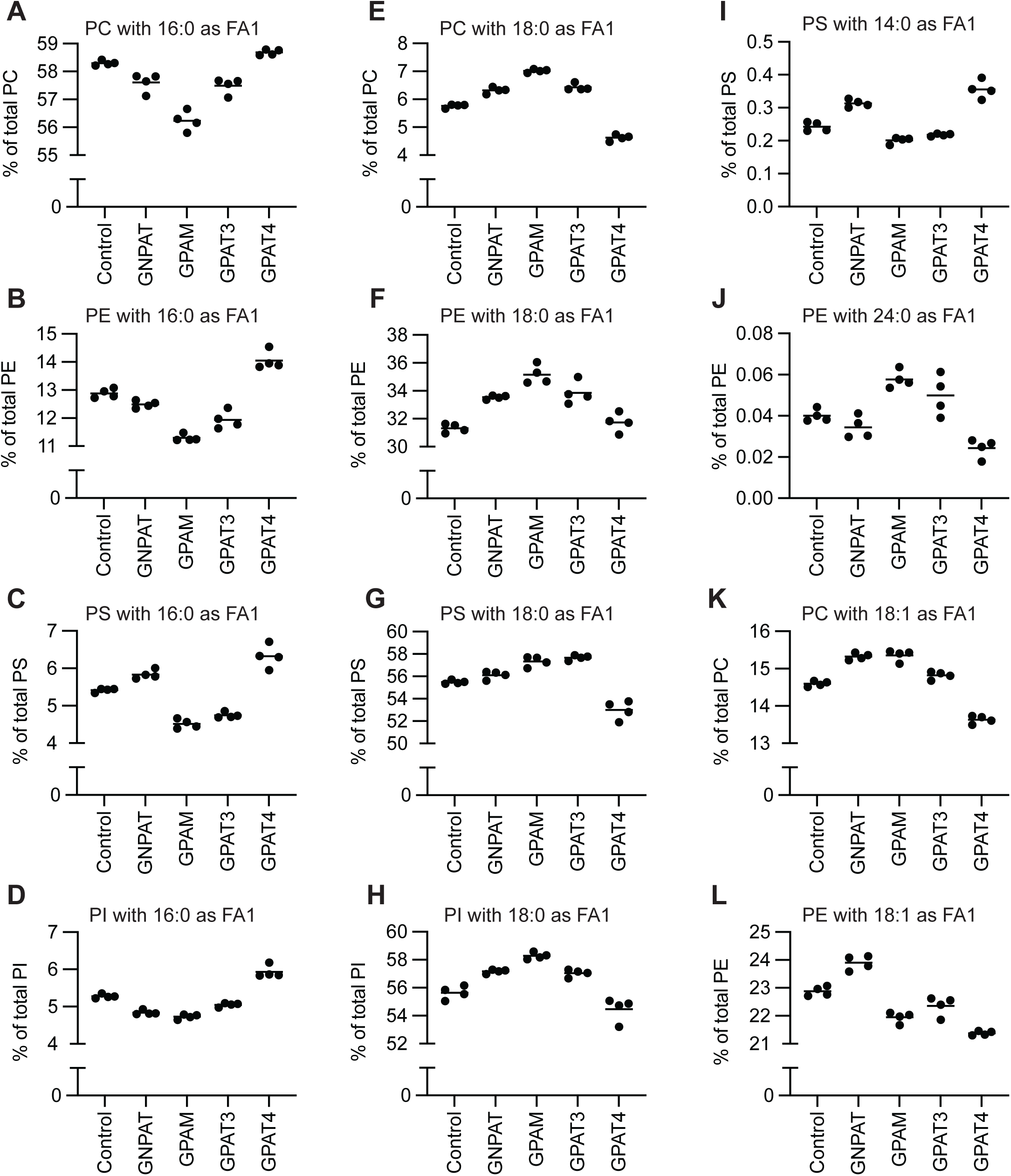
Analysis of acyl chain changes in GPAT/GNPAT mutants. (A-L) Levels of the indicated acyl chains constituting FA1 in individual GPL classes, in the indicated mutants. Results of statistical tests are provided in supplemental file 3.

Based on the loading analysis of PC2 described above, we investigated 15:0 changes in the mutants, and found decreases in GNPAT mutants (Figure 5A). We noticed that odd-chain GPLs often appear as double peaks in reverse-phase chromatography (Figure 5B). The early-eluting peaks, which we label here as peak 1, have retention times that are earlier than what is expected from the elution times of even-chain GPLs (Figure 5C). Thus, these early-eluting odd-chain GPLs probably contain acyl chains that are structurally divergent, which we speculate to be branched-chain. GNPAT mutants had decreases mainly in the first peaks (Figure 5BDE), suggesting that GNPAT incorporates atypical odd-chain acyl chains. Therefore, our analysis revealed that the balance between GPAM and GPAT4 affects saturated acyl chain length, while GPAT3 is more redundant and GNPAT contributes to atypical odd-chain acyl chain incorporation. The analysis also showed that GPATs/GNPAT single mutants do not have extreme acyl chain changes. This showed the functional redundancy between these enzymes, which we explored further.

**Figure 5.**
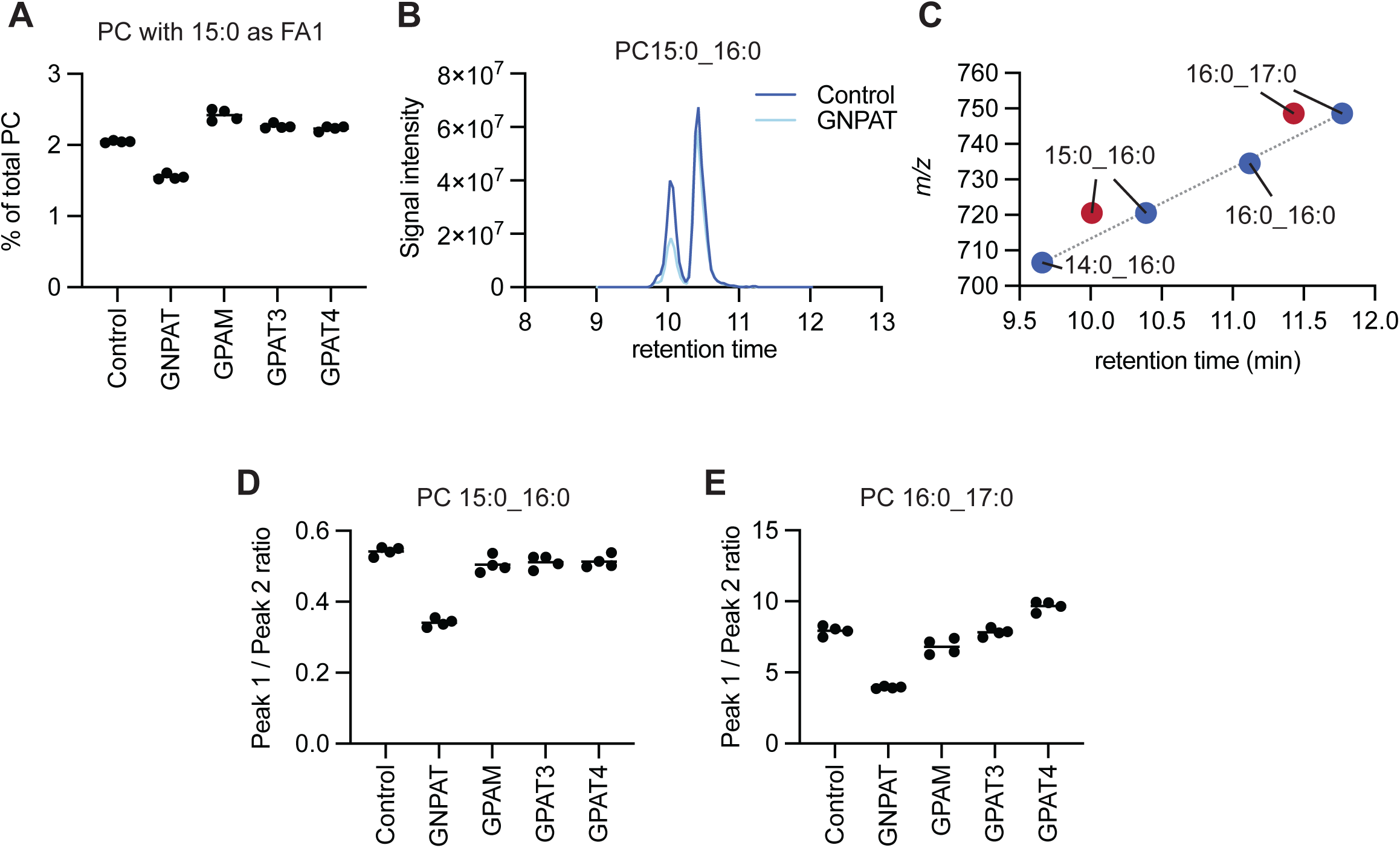
Changes in odd-chain acyl chains in GNPAT mutants. (A) Levels of 15:0 as FA1 of phosphatidylcholine. (B) Representative chromatograms of PC 15:0_16:0 during LC-MS analysis of lipids from control and GNPAT mutant cells. Note a reduction in the first peak. (C) Comparison of retention times between distinct PC species. Odd-chain PC have two peaks, with the first ones eluting early than the expected retention time. (DE) Ratio between the two peaks of odd-chain PC species, with peak 1 being the one eluting earlier. Results of statistical tests are provided in supplemental file 3.

### Redundancy of GPATs/GNPAT

We next investigated the extent of redundancy between GPATs and GNPAT. Inspired by the study showing a synthetic lethality of GNPAT and CHP1^8^, the activator of GPAT4, we targeted GPAT4 and GNPAT alone or in combination, while also testing the effect of targeting all GPATs and GNPAT expressed in HeLa cells (Figure 6A). Hereafter, multiplex mutants are named based on the targeted genes, with GPAM, GPAT3, GPAT4, and GNPAT being labeled as M, 3, 4, and N, respectively (for example, GPAT4/GNPAT-targeted cells are named 4N). After the drug selection used for mutant generation^15^, we found normal survival of 4N cells, while quadruple-targeted M34N cells barely survived (Figure 6A). Analysis of the obtained mutants showed nearly perfect mutation rates in GPAT4 and GNPAT in all cell lines (single mutants, 4N, and M34N), while GPAM and GPAT3 mutation rates were modest in M34N cells, when compared to single gene mutants (compare Figures 2A and 6B). In addition, we found a high abundance of in-frame 3 base deletions in GNPAT gene of M34N cells, which was in clear contrast to GNPAT single mutants (Figure 6C). This showed that the few surviving M34N were not fully knocked out for all targets. Rather, in-frame mutations of GNPAT, as well as incomplete mutations in GPAM and GPAT3, enabled the survival of a small population of targeted cells, while all (or at least the majority of) cells successfully mutated for all targets died during the generation of the cell line. This showed the possibility to assess synthetic lethality of genes by analyzing whether in-frame mutations are over-represented in mutants. We analyzed in-frame mutations in all targets of the established mutants, and found that 4N cells are not enriched in in-frame mutations, while M34N cells have high in-frame mutation rates in GNPAT, GPAM, and GPAT3 (Figure 6D). Thus, from this analysis we concluded that in HeLa cells, combined GPAT4 and GNPAT mutations do not affect survival, while mutations in all GPATs and GNPAT are lethal.

**Figure 6.**
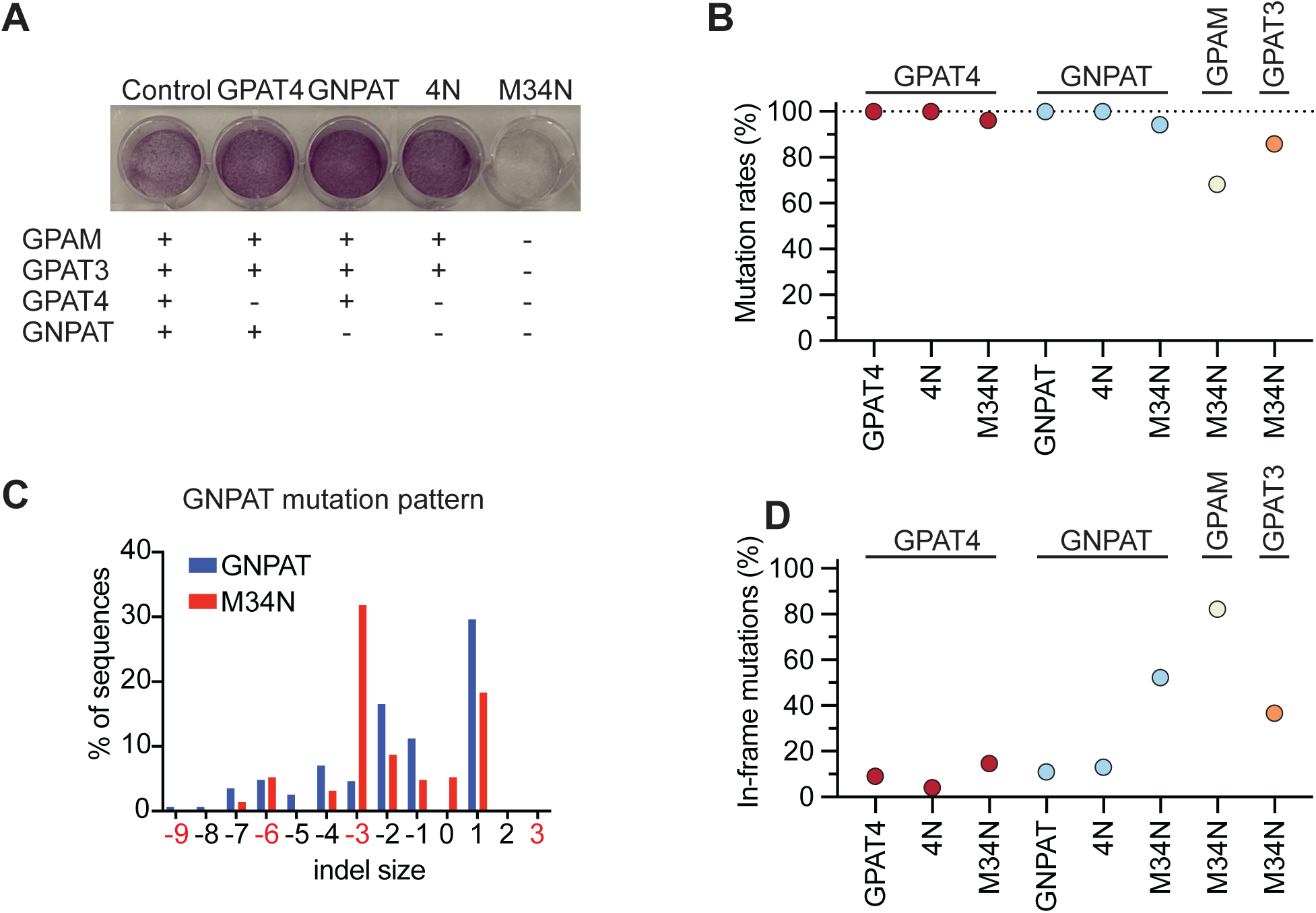
Analysis of redundancy between GPATs and GNPAT. (A) Survival analysis of the indicated mutants. Cells that survived after drug selection used for mutant generation were reseeded and stained with crystal violet. Genes labeled with a minus are those that were targeted. (B) Mutation rates of each target (shown above the plots) in individual mutant cells (shown under the X axis), as analyzed by TIDE. (C) Mutation patterns of GNPAT in a single mutant or M34N cells, as analyzed by TIDE. See the abundance of the -3 peak in M34N, illustrating an in-frame mutation of GNPAT. (D) Percentage of in-frame mutations obtained in each target (shown above the plots) in individual mutant cells (shown under the X axis). The results are from a single analysis.

We further repeated the experiment with more combinations of targeting, including all possible triple mutants (Figure 7A-C). We found that most triple mutants (M4N, M3N, M34) have no obvious lethality, with the exception of 34N cells, in which only GPAM is left (Figure 7A). Mutation rates were high in all the generated mutants (Figure 7B), but we barely obtained 34N cells, in which we found high in-frame mutation rates in GNPAT (Figure 7C). Nevertheless, in contrast to M34N cells, in-frame mutation rates were not increased for GPAT3 in 34N cells. In addition, in-frame rates of GNPAT mutation were lower in 34N than in M34N cells (Figure 7C). Altogether, our results suggested that combinatory mutations of GPATs and GNPAT are mostly tolerated, unless all of them are mutated. In addition, 34N mutants were likely to have defects in proliferation, suggesting that GPAM-mediated LPA synthesis is not sufficient for normal rates of cell growth. On the other hand, there was no evidence of lethality or proliferation defects in M34 cells, in which only GNPAT is untargeted (Figure 7A-C). This suggested that peroxisome-derived LPA synthesis is sufficient to support normal cell growth. We concluded that as compared to mitochondria, peroxisomes are a better source of LPA for GPL synthesis in ER. This establishes GNPAT as an important enzyme for the synthesis of GPLs, in addition to its traditionally known role in ether lipid synthesis.

**Figure 7.**
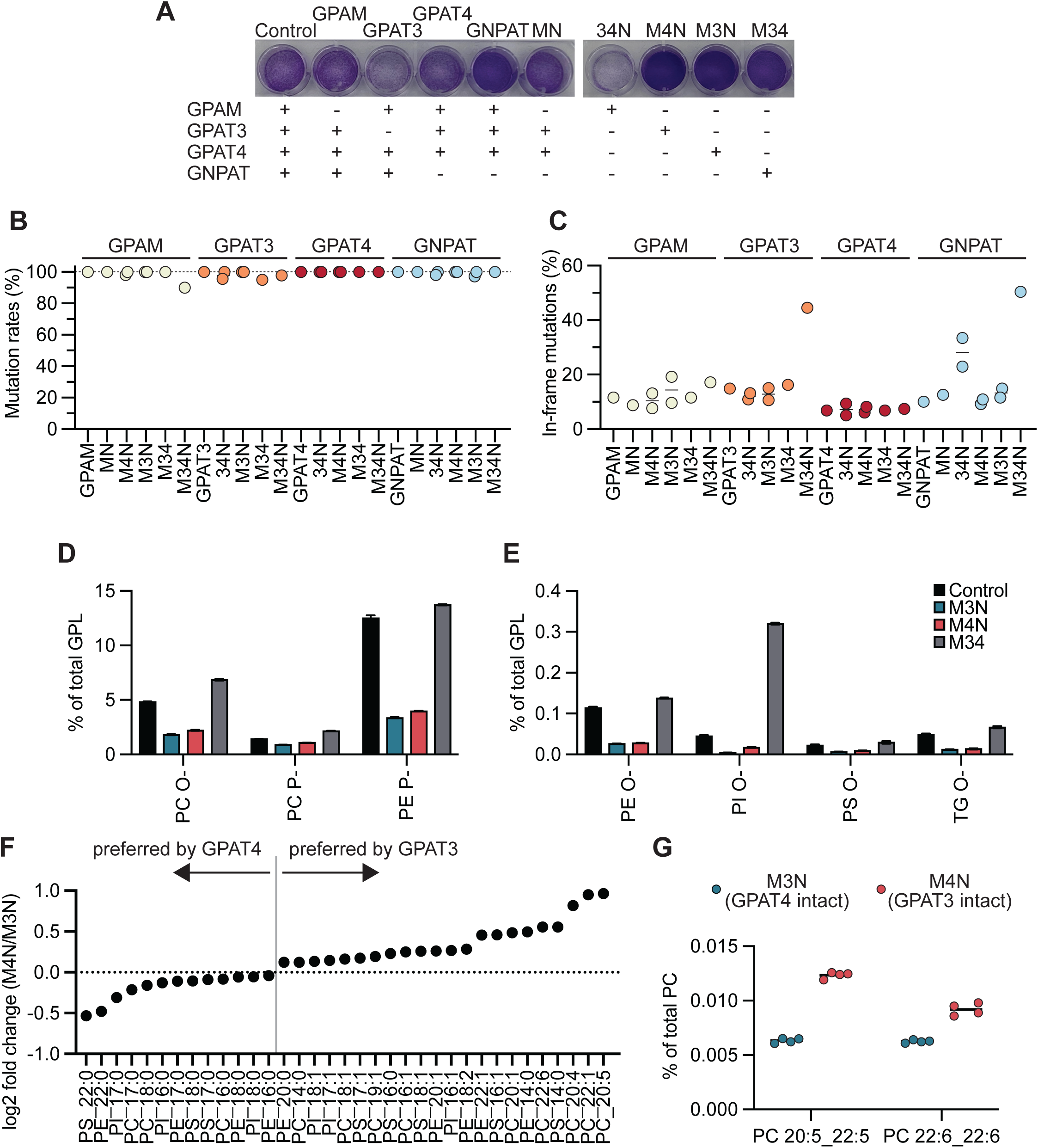
Lipid changes in multiplex GPAT/GNPAT mutants. (A) Survival analysis of the indicated mutants. (B) Mutation rates and (C) percentages of in-frame mutations obtained in the indicated mutant cells. See also Figure 6 legend. (DE) Changes in ether lipid species in the indicated mutants. Error bars are SEM (n=4). (F) Changes in acyl chains that constitute FA1 in M3N (expressing only GPAT4) and M4N mutants (expressing only GPAT3). (G) Comparison of di-polyunsaturated PC species in M3N and M4N mutants. Results of statistical tests are provided in supplemental file 3.

### Comparison of lipid compositions in triple mutants

We took advantage of the triple mutants that we established to further analyze the roles of GPATs and GNPAT in lipid regulation. M34 cells, with only GNPAT left, had increased ether lipid levels, which were relatively at moderate extents (Figure 7DE). This showed again that GNPAT alone sufficiently covers the synthesis of GPLs, including those without ether linkages. Ether PI species were drastically increased (7-fold) in M34 cells (Figure 7E), confirming the above observation that increased flux toward ether lipid synthesis is especially reflected in ether PI levels. Ether lipids were not decreased in 34N cells, which was consistent with the high in-frame mutation rates seen in GNPAT (data not shown). Thus, GNPAT in-frame mutations left some functional proteins, which probably contributed to cell proliferation defects that would happen if only GPAM was remaining.

Finally, we compared M3N and M4N cells, which would reveal the preferences of the two ER GPATs. We assumed that the role of GPAT3 would be clearer in this comparison, due to the lack of GPAM that had similar functions. By comparing acyl chains (only FA1 were analyzed, as was done for single mutants) that changed in the two mutants (pre-selected by false discovery rate-corrected multiple t-test analyses), we found that GPAT4 has a higher contribution on saturated acyl chain incorporation, while GPAT3 prefers unsaturated substrates when compared to GPAT4 (Figure 7F). The most drastic differences between the two enzymes were seen in polyunsaturated acyl-chains, which were better incorporated by GPAT3 (Figure 7F). Consistently, the levels of di-polyunsaturated PC species were largely different between M3N and M4N mutants. Thus, this additional analysis revealed that the balance between GPAT3 and GPAT4 affect the degree of unsaturation in acyl chains.

## 4. Discussion

### Regulation of ether lipids by GPATs and GNPAT

By analyzing combinatory mutant cells of GPATs and GNPAT, we established the regulatory roles of these enzymes on ether lipids. We found that ether lipids increase when the relative contribution of GNPAT on lipid synthesis becomes more important, as a consequence of GPAT mutations. Interestingly, this trend was especially pronounced for ether PI, which is a typically ignored, very minor GPL subclass. We speculate that additional regulatory mechanisms exist for the major ether lipids, alkyl PC and alkenyl PE, so that they do not over-accumulate in cells. In such scenario, even when GNPAT has extreme contribution, the flux of alkyl-LPA toward alkyl PC and alkenyl PE remains relatively controlled, thus leading only to moderate increases. This would lead to an over-accumulation of ether lipid precursors, such as alkyl phosphatidic acid, which can be converted into ether PI. Thus, our results suggest that ether PI can be a good marker of peroxisome-derived lipid synthesis. Ether lipids are thought to have important cellular functions, while being altered in various diseases including cancer^10,20^. Ether lipid deficiency leads to the severe genetic disease rhizomelic chondrodysplasia punctata^21^. Thus, understanding ether lipid regulation is critical, and our results suggest that GNPAT competition with GPATs is one of the regulatory factors. This suggests that GPAT inhibition could be a strategy to restore ether lipid levels.

### Regulation of acyl chains by GPATs and GNPAT

Our results suggest that GPATs and GNPAT have partially redundant regulatory roles on acyl chains. First, GPATs/GNPAT mutations lead acyl chain changes that are partial, showing that other enzymes cover the functions of the mutated enzyme(s). Thus, as far as the substrate are supplied, it is likely that most GPATs and GNPAT are able to synthesize LPA species with the acyl chains required for normal cell functions. Nevertheless, we found significant differences in the enzymes that would contribute to the fine tuning of membrane compositions. GPAM and GPAT4 have opposite effects on saturated acyl chain length, with C17 being the threshold. This threshold is interesting, as it is in the middle of the most commonly found *sn-*1 acyl chains, which are of C16 and C18 chain lengths. Thus, a precise regulation between the most abundant acyl chains is likely to have the highest biological relevance, which would explain why evolution led to a threshold of C17 between GPAM- and GPAT4-regulated acyl chains. Indeed, multiple evidence suggest that the ratio between C16 and C18 saturated acyl chains are important. For example, PS having oleic acid (18:1) at the *sn-*2 position interacts with cholesterol when its *sn-*1 acyl chain is stearic acid (18:0), but not palmitic acid (16:0)^22^. The enzyme LPGAT1 was shown to remodel GPLs, especially PE, to increase 18:0 levels at the *sn-*1 position, and its deficiency is partially lethal^23,24^. The existence of LPGAT1 as a regulator of 18:0 in PE might also explain why GPAT4 mutants had no changes in this acyl chain for PE, in contrast to PC, PI, and PS. The studies discussed here demonstrate the importance of proper *sn-*1 acyl chain regulation, and therefore the balance between GPAM and GPAT4 is likely to have important cellular roles.

We found that GNPAT has higher contribution than GPATs for atypical, possibly branched-chain, acyl chains. Branched-chain fatty acids are potentially beneficial for our health^25^, and thus a confirmation of GNPAT roles in their incorporation would help the clarification of their functions. We also found that GPAT3 and GPAT4 substantially differ in substrate preferences, with GPAT3 being more adapted for polyunsaturated acyl chain incorporation. We note that M4N cells, which have only GPAT3 intact, still maintain relatively normal acyl chain profiles. Thus, GPAT3 is sufficiently selective for acyl-CoA species that would be needed to maintain typical 1-saturated/2-unsaturated asymmetric GPLs. While polyunsaturated acyl chains are typically minor at the *sn-*1 position, retinal lipids are rich in di-polyunsaturated species^26^. It would be interesting to investigate whether GPAT3 or other GPATs contribute to the synthesis of such lipids.

### Redundancies in GPATs and GNPAT from distinct organelles

Our data suggest that LPA synthesis in mitochondria does not support bulk lipid synthesis required for efficient cell growth, while LPA synthesis initiated in peroxisomes does. This result was unexpected, since the ER has contacts with both mitochondria and peroxisomes, and ER-mitochondria contacts have established roles in lipid exchange^27^. This raises the possibility that LPA is not a good substrate for lipid transporters working between the ER and mitochondria. This would explain the necessity of having GPATs in both ER and mitochondria. On the other hand, the sufficiency of peroxisome-derived lipid synthesis to support cell growth is expectable, as cell normally use the shuttling of alkyl-LPA (or alkyl-DHAP) from peroxisomes to the ER for ether lipid synthesis. Thus, cells have an established system to transport LPA between the organelles, thus explaining the sufficiency of GNPAT to initiate sufficient lipid synthesis for cell growth.

## Conclusion

Our study revealed the multifaceted functions of GPATs and GNPAT, in the regulation of ether lipids, acyl chains, and bulk lipid synthesis. Knockout mouse studies revealed the functions of these enzymes^12^, but the molecular mechanisms of lipid functions affected in these mice have been typically difficult to investigate. Our study contributes to a better understanding about the metabolic functions of these enzymes, and thus will help the establishment of hypotheses when investigate the knockout mice in the future. Lipidomics became now a major tool in biology and medical research^28^, and its ability to detect individual lipid species makes it critical to study how lipid profiles are fine tuned. Our study unveils how the first step of de novo lipid synthesis contributes to lipid compositions, and similar analyses of other pathways would lead to a complete understanding of lipidome regulation.

## Supporting information

Supplemental File 1, primer list

Supplemental File 2, lipidomics data

Supplemental file 3, statistics

## Acknowledgements

We are grateful to the members of the Harayama laboratory for their support and discussion. We thank Bruno Antonny (Institut de Pharmacologie Moléculaire et Cellulaire) and his lab members for their support. We thank Hiroshi Tsugawa, Yuki Matsuzawa, and Manami Takeuchi (Tokyo University of Agriculture and Technology) for software development and assistance in lipidomics analyses. We thank Nathalie Leroudier (Institut de Pharmacologie Moléculaire et Cellulaire) for Sanger sequencing analysis. We thank Daisuke Hishikawa (Nippon Medical School) and Keisuke Yanagida (National Center for Global Health and Medicine) for critical reading of the manuscript. T.H. was supported by the French Government (National Research Agency, ANR) through the “Investments for the Future” programs LABEX SIGNALIFE ANR-11-LABX-0028 and IDEX UCAJedi ANR-15-IDEX-01, the ATIP-Avenir program (CNRS/Inserm), and by the Global Innovation Research funds of Tokyo University of Agriculture and Technology.

## 5. Authors contributions

SS designed the research, generated mutant cell lines, and did lipidomics data analysis. HQ generated mutant cell lines, performed survival analysis, and prepared lipidomics samples. GPJ contributed to unpublished data required for correct annotations of lipidomics peaks. LF and DD did lipid extraction and LC-MS analyses for lipidomics. TH designed the research, did lipidomics data analysis, statistics, and wrote the manuscript. All authors read and edited the manuscript.

---

**Figure S1.**
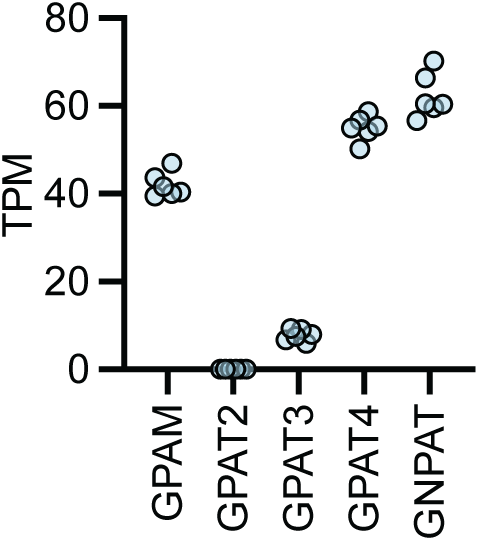
Expression of target genes in Hela cells. The expression of each target gene, as obtained from published RNA-seq data^29^.

**Figure S2.**
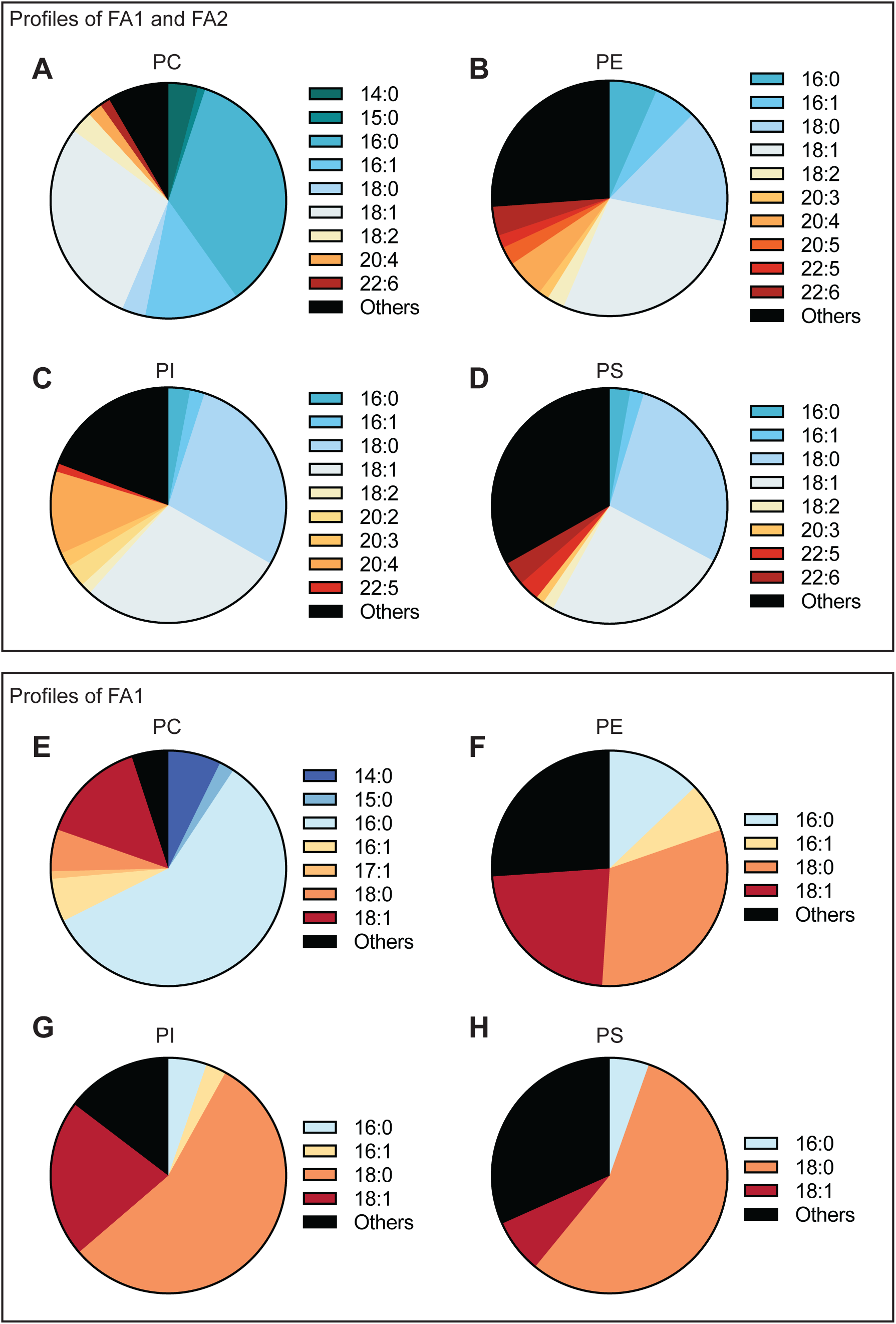
Acyl chain profiles of control cells. (A-D) Percentage of acyl chains constituting FA1 and FA2 in the indicated GPLs of control cells. Only species with >1% abundance are shown, while “Others” illustrate the minor species and the species for which unequivocal acyl chain annotation could not be done by LC-MS. (E-H) Same as (A-D) but with only FA1 analyzed.

## References

1. Harayama, T., and Riezman, H. (2018). Understanding the diversity of membrane lipid composition. Nature Reviews Molecular Cell Biology 19, 281–296. 10.1038/nrm.2017.138

2. Harayama, T. (2023). Metabolic bias: Lipid structures as determinants of their metabolic fates. Biochimie 215, 34–41. 10.1016/j.biochi.2023.09.019

3. Manni, M.M., Tiberti, M.L., Pagnotta, S., Barelli, H., Gautier, R., and Antonny, B. (2018). Acyl chain asymmetry and polyunsaturation of brain phospholipids facilitate membrane vesiculation without leakage. eLife 7, e34394. 10.7554/eLife.34394

4. Ariyama, H., Kono, N., Matsuda, S., Inoue, T., and Arai, H. (2010). Decrease in Membrane Phospholipid Unsaturation Induces Unfolded Protein Response. Journal of Biological Chemistry 285, 22027–22035. 10.1074/jbc.M110.126870

5. Volmer, R., van der Ploeg, K., and Ron, D. (2013). Membrane lipid saturation activates endoplasmic reticulum unfolded protein response transducers through their transmembrane domains. Proceedings of the National Academy of Sciences 110, 4628–4633. 10.1073/pnas.1217611110

6. Levental, K.R., Malmberg, E., Symons, J.L., Fan, Y.-Y., Chapkin, R.S., Ernst, R., and Levental, I. (2020). Lipidomic and biophysical homeostasis of mammalian membranes counteracts dietary lipid perturbations to maintain cellular fitness. Nature Communications 11, 1339. 10.1038/s41467-020-15203-1

7. Valentine, W.J., Shimizu, T., and Shindou, H. (2023). Lysophospholipid acyltransferases orchestrate the compositional diversity of phospholipids. Biochimie 215, 24–33. 10.1016/j.biochi.2023.08.012

8. Zhu, X.G., Nicholson Puthenveedu, S., Shen, Y., La, K., Ozlu, C., Wang, T., Klompstra, D., Gultekin, Y., Chi, J., Fidelin, J., et al. (2019). CHP1 Regulates Compartmentalized Glycerolipid Synthesis by Activating GPAT4. Molecular Cell 74, 45–58.e47. 10.1016/j.molcel.2019.01.037

9. Wendel, A.A., Lewin, T.M., and Coleman, R.A. (2009). Glycerol-3-phosphate acyltransferases: Rate limiting enzymes of triacylglycerol biosynthesis. Biochimica et Biophysica Acta (BBA) - Molecular and Cell Biology of Lipids 1791, 501–506. 10.1016/j.bbalip.2008.10.010

10. Jiménez-Rojo, N., and Riezman, H. (2019). On the road to unraveling the molecular functions of ether lipids. FEBS Letters 593, 2378–2389. 10.1002/1873-3468.13465

11. Ohba, Y., Sakuragi, T., Kage-Nakadai, E., Tomioka, N.H., Kono, N., Imae, R., Inoue, A., Aoki, J., Ishihara, N., Inoue, T., et al. (2013). Mitochondria-type GPAT is required for mitochondrial fusion. The EMBO Journal 32, 1265–1279. 10.1038/emboj.2013.77

12. Huang, Y., Hu, K., Lin, S., and Lin, X. (2022). Glycerol-3-phosphate acyltransferases and metabolic syndrome: recent advances and future perspectives. Expert Reviews in Molecular Medicine 24, e30. 10.1017/erm.2022.23

13. Lewin, T.M., de Jong, H., Schwerbrock, N.J.M., Hammond, L.E., Watkins, S.M., Combs, T.P., and Coleman, R.A. (2008). Mice deficient in mitochondrial glycerol-3-phosphate acyltransferase-1 have diminished myocardial triacylglycerol accumulation during lipogenic diet and altered phospholipid fatty acid composition. Biochimica et Biophysica Acta (BBA) - Molecular and Cell Biology of Lipids 1781, 352–358. 10.1016/j.bbalip.2008.05.001

14. Shiozaki, Y., Miyazaki-Anzai, S., Okamura, K., Keenan, A.L., Masuda, M., and Miyazaki, M. (2020). GPAT4-Generated Saturated LPAs Induce Lipotoxicity through Inhibition of Autophagy by Abnormal Formation of Omegasomes. iScience 23, 101105. 10.1016/j.isci.2020.101105

15. Harayama, T., Hashidate-Yoshida, T., Aguilera-Romero, A., Hamano, F., Morimoto, R., Shimizu, T., and Riezman, H. (2020). Establishment of a highly efficient gene disruption strategy to analyze and manipulate lipid co-regulatory networks. bioRxiv 2020.11.24.395632. 10.1101/2020.11.24.395632

16. Brinkman, E.K., Kousholt, A.N., Harmsen, T., Leemans, C., Chen, T., Jonkers, J., and van Steensel, B. (2018). Easy quantification of template-directed CRISPR/Cas9 editing. Nucleic Acids Research 46, e58. 10.1093/nar/gky164

17. Tokiyoshi, K., Matsuzawa, Y., Takahashi, M., Takeda, H., Hasegawa, M., Miyamoto, J., and Tsugawa, H. (2023). Using data-dependent and independent hybrid acquisitions for fast liquid chromatography-based untargeted lipidomics. bioRxiv, 2023.2010.2012.562117. 10.1101/2023.10.12.562117

18. Koch, J., Lackner, K., Wohlfarter, Y., Sailer, S., Zschocke, J., Werner, E.R., Watschinger, K., and Keller, M.A. (2020). Unequivocal Mapping of Molecular Ether Lipid Species by LC–MS/MS in Plasmalogen-Deficient Mice. Analytical Chemistry 92, 11268–11276. 10.1021/acs.analchem.0c01933

19. Dorninger, F., Brodde, A., Braverman, N.E., Moser, A.B., Just, W.W., Forss-Petter, S., Brügger, B., and Berger, J. (2015). Homeostasis of phospholipids — The level of phosphatidylethanolamine tightly adapts to changes in ethanolamine plasmalogens. Biochimica et Biophysica Acta (BBA) - Molecular and Cell Biology of Lipids 1851, 117–128. 10.1016/j.bbalip.2014.11.005

20. Papin, M., Bouchet, A.M., Chantôme, A., and Vandier, C. (2023). Ether-lipids and cellular signaling: A differential role of alkyl- and alkenyl-ether-lipids? Biochimie 215, 50–59. 10.1016/j.biochi.2023.09.004

21. Koch, J., Watschinger, K., Werner, E.R., and Keller, M.A. (2022). Tricky Isomers—The Evolution of Analytical Strategies to Characterize Plasmalogens and Plasmanyl Ether Lipids. Frontiers in Cell and Developmental Biology 10, 864716. 10.3389/fcell.2022.864716

22. Maekawa, M., and Fairn, G.D. (2015). Complementary probes reveal that phosphatidylserine is required for the proper transbilayer distribution of cholesterol. Journal of Cell Science 128, 1422–1433. 10.1242/jcs.164715

23. Sato, T., Umebayashi, S., Senoo, N., Akahori, T., Ichida, H., Miyoshi, N., Yoshida, T., Sugiura, Y., Goto-Inoue, N., Kawana, H., et al. (2023). LPGAT1/LPLAT7 regulates acyl chain profiles at the sn-1 position of phospholipids in murine skeletal muscles. Journal of Biological Chemistry 299, 104848. 10.1016/j.jbc.2023.104848

24. Xu, Y., Miller, P.C., Phoon, C.K.L., Ren, M., Nargis, T., Rajan, S., Hussain, M.M., and Schlame, M. (2022). LPGAT1 controls the stearate/palmitate ratio of phosphatidylethanolamine and phosphatidylcholine in sn-1 specific remodeling. Journal of Biological Chemistry 298, 101685. 10.1016/j.jbc.2022.101685

25. Taormina, V.M., Unger, A.L., Schiksnis, M.R., Torres-Gonzalez, M., and Kraft, J. (2020). Branched-Chain Fatty Acids—An Underexplored Class of Dairy-Derived Fatty Acids. Nutrients 12, 2875. 10.3390/nu12092875

26. Rice, D.S., Calandria, J.M., Gordon, W.C., Jun, B., Zhou, Y., Gelfman, C.M., Li, S., Jin, M., Knott, E.J., Chang, B., et al. (2015). Adiponectin receptor 1 conserves docosahexaenoic acid and promotes photoreceptor cell survival. Nature Communications 6, 6228. 10.1038/ncomms7228

27. Wu, H., Carvalho, P., and Voeltz, G.K. (2018). Here, there, and everywhere: The importance of ER membrane contact sites. Science 361, eaan5835. 10.1126/science.aan5835

28. Ni, Z., Wölk, M., Jukes, G., Mendivelso Espinosa, K., Ahrends, R., Aimo, L., Alvarez-Jarreta, J., Andrews, S., Andrews, R., Bridge, A., et al. (2022). Guiding the choice of informatics software and tools for lipidomics research applications. Nature Methods 20, 193–204. 10.1038/s41592-022-01710-0

29. Scott, C.C., Vossio, S., Rougemont, J., and Gruenberg, J. (2018). TFAP2 transcription factors are regulators of lipid droplet biogenesis. eLife 7, e36330. 10.7554/eLife.36330

